# Mitochondrial uncoupler BAM15 improves skeletal muscle function and mitochondrial respiration in Sarcopenia

**DOI:** 10.1101/2025.10.30.685477

**Authors:** Dean G. Campelj, Ashleigh M. Philp, Elya J. Ritenis, Camila S. Padilha, Isabelle Alldritt, James Sligar, Tabitha Cree, Stephanie J. Alexopoulos, Webster. L Santos, Sophie Joanisse, Paul M. Coen, Kyle L. Hoehn, Andrew Philp

**Affiliations:** Centre for Healthy Ageing, Centenary Institute, Sydney, Australia; Faculty of Medicine and Health, Charles Perkins Centre, University of Sydney, Sydney, Australia; School of Clinical Medicine, UNSW Medicine and Health, St Vincent’s Healthcare clinical campus, University of New South Wales, Sydney, Australia; Centre for Inflammation, Centenary Institute, and Faculty of Science, University of Technology Sydney, Sydney, Australia; Department of Health Sciences and Biostatistics, Swinburne University of Technology, Melbourne, Australia; School of Sport, Exercise and Rehabilitation Sciences, University of Technology Sydney, Sydney, Australia; Murdoch Children’s Research Institute, The Royal Children’s Hospital, Melbourne, Australia; reNEW, Novo Nordisk Foundation Centre for Stem Cell Medicine, Melbourne, Australia; School of Biotechnology and Biomolecular Sciences, University of New South Wales, Sydney, Australia; Department of Chemistry and Virginia Tech Center for Drug Discovery, Virginia Tech, Blacksburg, USA; School of Life Sciences, University of Nottingham, Nottingham, United Kingdom; Translational Research Institute for Metabolism and Diabetes, AdventHealth, Orlando, USA

**Keywords:** Sarcopenia, mitochondrial uncouplers, skeletal muscle dysfunction, mitochondrial respiration, oxidative stress

## Abstract

**Background:** Ageing is accompanied by progressive declines in skeletal muscle mass and strength, culminating in sarcopenia, a condition that contributes to frailty, multimorbidity, and mortality. Age-related changes to mitochondria lead to oxidative damage and dysfunction and are proposed to occur early in the trajectory of sarcopenia, supporting the candidacy of mitochondrial-protective therapies. Here, we test the efficacy of mitochondrial uncoupler BAM15 in age-dependent sarcopenic mouse models.

**Methods:** Male and female MitoQC mice aged 24 months received either standard chow or chow supplemented with BAM15 (0.033% mg/g) ad libitum for eight weeks (n=13–14/group). Young (3-month-old) mice served as reference controls (n=8/group). Muscle mitochondrial respiration was assessed in permeabilized fib res, and contractile function was measured in isolated extensor digitorum longus and soleus muscles. Mitophagy was quantified by immunofluorescence confocal microscopy. Data were analyzed using one-or two-way ANOVA followed by Dunnett’s or Bonferroni’s multiple comparison tests.

**Results:** Aged male and female mice exhibited reduced gastrocnemius muscle mass relative to body mass compared with young controls (p<0.05; ∼18% and ∼32% loss, respectively). BAM15 did not alter muscle size but reversed the age-related loss of contractile function in EDL muscles, to that of the young reference controls in both sexes (p<0.05; ∼33% in males, ∼16% in females). In male mice, BAM15 improved mitochondrial efficiency, evidenced by restoration of Complex I–linked respiration and decreased proton leak (∼52% improvement; p<0.05), and normalized protein levels of oxidative stress marker 4 -HNE, without changes in mitophagy or mitochondrial content. In females, BAM15 did not improve mitochondrial parameters, which may be, in part, due to aged female muscle exhibiting unchanged Complex I leak and 4-HNE protein abundance, alongside lower complex I subunit (NDUFB8) protein abundance.

**Conclusions:** BAM15 improved skeletal muscle mitochondrial efficiency and contractile function in aged male mice, supporting the potential of mitochondrial uncoupling as a therapeutic strategy for sarcopenia.

## Introduction

Sarcopenia is the clinical term for age-related skeletal muscle decline, characterized by loss of muscle mass, weakness and reduced physical capacity [1]. This culminates into frailty conditions, which are associated with increased risk of falls and fractures, hospitalization, health care costs, co-morbidities and mortality [2]. To date, therapeutic interventions for the treatment of sarcopenia are limited, and those proposed often fail to show functional improvements in physical capacity [3]. Therefore, novel strategies to protect against age-related muscle decline are warranted.

Mitochondrial dysfunction is a hallmark of ageing with several mechanisms proposed [4]. Accordingly, mitochondrial uncoupler compounds are being explored as potential interventions to delay and prevent sarcopenia progression [5-7]. Mitochondria provide cellular energy for biological processes, primarily through electron transport chain (ETC) oxidative phosphorylation. This process pumps protons from the mitochondrial matrix into the intermembrane space to create the protonmotive force(PMF) that couples substrate oxidation to ATP synthesis [8]. In ageing, compromised ETC functional capacity leads to elevated PMFand increased reactive oxygen species (ROS) production. Mitochondrial uncoupler compounds lower the PMF and decreases ROS production by transporting protons from the mitochondrial intermembrane space into the mitochondrial matrix [5, 6].

Here, we evaluated the mitochondrial uncoupler BAM15 as a therapeutic for sarcopenia. BAM15 is a mitochondrial-targeted protonophore ((2-fluorophenyl){6-[(2-fluorophenyl)amino](1,2,5-oxadiazolo[3,4-e]pyrazin-5-yl)}amine) that has been shown to prevent diet-induced obesity, insulin resistance [9] and skeletal muscle decline in sarcopenic-obesity mouse models [10]. Unlike traditional uncouplers such as carbonyl cyanide p-trifluoromethoxyphenylhydrazone (FCCP) [11], BAM15 does not induce mitochondrial membrane depolarization or damage and shows excellent bioavailability in skeletal muscle following oral delivery [9]. We hypothesised that short-term, mild mitochondrial uncoupling via BAM15 supplementation, would preserve skeletal muscle mitochondrial function and thereby protect against age-associated muscle decline.

## Methods

### Experimental protocols and treatments

#### Mouse cohort

All animal experiments were conducted in compliance with Garvan Ethics Committee guidelines (21/20 protocols) and in accordance with the Australian code of practice for the care and use of animals for scientific purposes. Mitophagy reporter mice (MitoQC) [12] were kindly provided by Professor Ian Ganley (University of Dundee, UK). The MitoQC construct is a constitutive knockin of tandem fluorophore tag mCherry and GFP on outer mitochondrial membrane protein FIS1^101–153^. MitoQC colony breeding and maintenance were completed at Australian Bioresources (Moss Vale, NSW, Australia) and genotyping was performed by Garvan Molecular Genetics (Darlinghurst, NSW, Australia). Mice were transferred to the Biological Testing Facility at Garvan Institute of Medical Research and were housed on a 12-hour light/dark cycle with ad libitum access to standard rodent chow and water. Once MitoQC mice were assigned to the aging colony and became 12-months of age they received bi-weekly health-checks and regular welfare monitoring by researchers and animal technicians until they reached the desired experimental age, i.e., 24-months-old.

#### Study design

At 24 months of age, male and female MitoQC mice were randomly allocated to an 8-week diet-intervention of standard chow (26M CHOW) or BAM15 supplemented chow (26M BAM15; (0.033% (mg/g)), with *n*=14/group. The age range of 24–26 months in the C57BL/6J mouse strain corresponds with established sarcopenia model classifications, characterized by loss of muscle mass and function [13]. These mice received daily welfare monitoring alongside tri-weekly recording of body weight and food intake. The dosing strategy for BAM15 was adapted from our previous work in C57BL/6J mice [9]. 1 male 26M CHOW and 1 female 26M BAM15 supplemented mice were euthanised during treatment due to reaching ethical humane endpoints and were excluded from analysis. Untreated 3-month-old male and female MitoQC mice (3M CON: *n*=8/group) were used as a healthy reference control group. Power calculations were based on our previous publication [14], with a minimum of n=8 aged mice per group was estimated to be sufficient to detect a change in muscle strength with age. To accommodate age-related attrition, group sizes were increased by ∼50%, consistent with reported survival rates from the Jackson Laboratory of C57BL/6 mice by 28 months of age.

#### BAM15 Diet Preparation

BAM15 was prepared as previously described [15]. Briefly, standard chow was first powdered using a food processor and used for all treatment groups. Powdered chow was then mixed with the correct concentration of BAM15 and compressed into pellets using a small pellet mill (Gemco ZLSP-120B). For the standard chow diet, powdered diet was pressed into pellets without adding any drug.

#### Tissue collection and processing

At study end, mice were anaesthetized using isoflurane (5% induction and 2-3% maintenance). Muscles and organs were surgically removed, weighed and collected via snap-freezing in liquid nitrogen or immersive fixation (3.7% PFA in 200 mM HEPES, pH 7). Exceptions include: micro-surgical harvesting of EDL and soleus (SOL) muscles for contractility testing and collection of one gastrocnemius (GTN) muscle per mouse which was immediately incubated in ice-cold BIOPS buffer post-harvest (2.77 mM CaK2EGTA, 7.23 mM K2EGTA, 5.77 mM Na2ATP, 6.56 mM MgCl2-6H2O, 20 mM taurine, 15 mM Na2-phosphocreatine, 20 mM imidazole, 0.5 mM dithiothreitol, 50 mM MES hydrate, pH 7.1). Euthanasia was confirmed following cardiac removal.

#### Skeletal muscle contractile function

*Ex vivo* electrophysical testing of skeletal muscle contractility was conducted on EDL and SOL muscles as previously described [16]. Muscles were tied at both tendons with 4.0 surgical silk thread before being excised from the hindlimb and placed into individual organ baths of a Myodynamics Muscle Strip Myograph System (DMT). Each organ bath was filled with a modified Krebs-Henseleit solution (NaCl 118 mM, MgSO4·7H2O 1 mM, KCl 4.75 mM, KH2PO4 1 mM, CaCl2 2.5 mM, NaHCO3 24 mM and glucose 11 mM; pH 7.4), bubbled with carbogen (5% CO2 in O2) and maintained at 30°C. The optimal length of each muscle was established through the delivery of sequential twitch contractions, while incrementally stretching each muscle, until the point where the greatest twitch force produced was determined and the resting length was measured via the micromanipulator. Next, a force–frequency protocol was performed utilizing tetanic stimulations with a square wave of 0.2ms pulses, and a pulse train duration of 350ms and 500ms for the EDL and SOL, respectively. EDL and SOL muscles were stimulated at an incremental range of frequencies from 10-150 Hz, with a 3-min rest in between each stimulation. Specific force was calculated using equations as previously described [17]. To investigate muscle resilience, muscles underwent a three-minute fatigue protocol, with EDL muscles stimulated at 100Hz every 4s, and, SOL muscles stimulated at 80 HZ every 2s. Recovery was also assessed, with tetanic stimulations occurring up to 15 mins post-fatigue protocol. Surgical silk was removed and muscles were blotted before tendon-free dry mass was recorded. All data were collected and annotated using LabChart Pro version 8.0 software (ADInstruments).

#### High-resolution respirometry

*Ex vivo* mitochondrial respiration was assessed in GTN muscles by measuring oxygen consumption using an Oxygraph-2k (OROBOROS Instruments) as previously published [18]. GTN muscles were mechanically separated into 3-5mg fibre bundles before being permeabilized in 50 µg/µL of saponin in BIOPS buffer for 30 mins at 4 °C. Fibre bundles were weighed and loaded into Oxygraph-2k chambers calibrated and pre-loaded with 2 mL of MiRO5 buffer (0.5 mM EGTA, 3 mM MgCl2-6H2O, 60 mM lactobionic acid, 20 mM taurine, 10 mM KH2PO4, 20 mM HEPES, 110 mM D-sucrose, 1 g/l bovine serum albumin, pH 7.1) at 37 °C. Oxygen concentration was maintained between 150-220 µM throughout the SUIT assay. First, pyruvate (10 mM) and malate (2 mM) were injected to stimulate respiration via ETC cycling at Complex I. ADP was then injected at incremental concentrations of 0.1 mM, 0.175 mM, 0.25 mM, 0.5 mM, 1 mM, 2 mM, 4 mM, and 6 mM, to generate a dose-response curve that depicts ADP-dependent oxidative phosphorylation. ADP sensitivity can then be determined through transforming data with the Michaelis–Menten enzyme kinetics fitting model using Prism (GraphPad Software, Inc.). Next, substrates glutamate (10 mM) and succinate (10 mM) were injected to assess phosphorylating respiration, followed by mitochondrial uncoupler Carbonyl cyanide m-chlorophenyl hydrazone (CCCP – 0.5 µM) to assess maximal ETC respiration. Lastly, Antimycin A (2.5 µM) was injected as a Complex III inhibitor, which ceases mitochondrial respiration and allows for the detection of non-mitochondrial respiration.

#### Mitophagy assessment via immunofluorescence

Quadriceps muscles were taken out of PFA 24-48 hours post fixation and placed in cryopreservation solution (30% sucrose in 1X PBS, pH 7) at 4 °C. Muscles were then placed in tissue moulds and embedded in optimal cutting temperature (OCT) compound (Sakura). Longitudinal cryosections were cut at 10 µm with a NX50 Cryostar Epredia Cryostat and mounted in ProLong Diamond Antifade (Thermofisher). Images were acquired using a Nikon C2-confocal microscope equipped with a 100X oil immersion objective. Image analysis was performed on ImageJ using semi-automatic plugin *mito-QC Counter* [18] to quantify mitolysosomes by red punctate counting, measuring mitolysosome size and mean channel intensities. Images were pre-processed by applying a Gaussian blur filter with a radius of 1-1.5μm and regions of interest drawn around muscle fibres for relative normalisation. Quantification of mitophagy was performed in 15-17 fibres per section.

#### Muscle lysate preparation and immunoblotting

Snap-frozen GTN muscles were prepared as previously published [18]. Briefly, GTN muscles were powdered on dry ice using a Cellcrusher tissue pulverizer (Cellcrusher Ltd) and homogenised via shaking in a Precellys Evolution Touch (Bertin Technologies) at 8,500 rpm for 2x 30 sec cycles in ice-cold sucrose lysis buffer (50 mM tris pH 7.5; 270 mM sucrose; 1 mM EDTA; 1 mM EGTA; 1% triton X-100; 50 mM sodium fluoride (NaF); 1 mM sodium orthovanadate (Na3VO4); 5 mM disodium pyrophosphate (Na2H2P2O7)) supplemented with fresh cOmplete™ EDTA free protease inhibitor cocktail (Roche) and phosphatase inhibitor cocktail (Sigma-Aldrich). Muscle homogenates were then centrifuged for 20 min at 14,800 rpm at 4°C to remove insoluble material and produce muscle lysates. Protein concentrations were determined using the RC/DC protein assay as per manufacturer’s instructions (Bio-Rad). Samples were prepared by mixing equivalent protein amounts of muscle lysates (30ug) in 4× Laemmli buffer (Bio-Rad) reduced in β-mercaptoethanol and boiled for 5 min at 95 °C. The antibody that required an exception to this protocol was the Total OXPHOS Rodent Antibody Cocktail, where samples remained unboiled per manufacturer’s instructions. Samples were separated by SDS-PAGE on 4-15% gradient gels (Bio-Rad) at 100 V and transferred onto nitrocellulose membranes (Bio-Rad) using a wet transfer system at 100 V for 60 min. Membranes were then stained in Ponceau S(Sigma-Aldrich) to check for even protein loading and transfer. Membranes were then blocked in 5% non-fat milk powder milk in tris-buffered saline with 0.1% Tween 20 (TBST) for 60 min followed by an overnight incubation at 4 °C with primary antibody dissolved in TBST containing either 3% BSA or non-fat milk powder. Primary antibodies probed include: Total OXPHOS Rodent Antibody Cocktail (1:1,000; ab110413; Abcam), Anti-NDUFB8 (1:1,000, ab192878, Abcam), Citrate Synthase (1:2,500, #14309, CST) and 4 Hydroxynonenal (1:1,000, ab46545, Abcam). After overnight incubation, membranes were washed 3 separate times for 5 min each in TBST and then probed with a horseradish peroxidase-conjugated secondary antibody (1:10,000; anti-rabbit IgG or anti-mouse IgG; CST) in TBST for 1 hour at room temperature. Following another set of 3 separate 10-min washes in TBST, images were developed using ECL reagent (Millipore), captured using the ChemiDoc Touch(Bio-Rad) and quantified using Image Labsoftware (Bio-Rad).

#### Statistics

Data are presented as Mean ± standard deviation and were analysed with GraphPad Prism v10. Normality of the data was assessed by Shapiro-Wilk tests. For parametric data, differences between experimental groups were determined by One-way ANOVA and Dunnet’s multiple comparisons testing or Two-way ANOVA and Bonferroni’s multiple comparisons testing. To detect significant differences in non-parametric data, a Kruskal-Wallis testing was performed followed by Dunn’s multiple comparisons testing. Significance level was set at *p*<0.05.

## Results

### BAM15 supplementation does not impact body and muscle mass in aged mice

To evaluate the potential therapeutic efficacy of BAM15 in sarcopenia, we supplemented 24-month-old male and female MitoQC mice for 8 weeks with either standard chow or 0.033% admixed BAM15 (Figure 1A). Anthropometric endpoint measures indicated that aged male and female mice had a sarcopenia-like phenotype with increased body mass (∼26% & ∼39%, for males and females, respectively), and reduced quadricep mass (∼22% & ∼17%, for males and females, respectively) and relative body to muscle mass ratios (∼22% & ∼35%, for males and females, respectively), compared to 3-month-old healthy controls (Figure 1B-1D). BAM15 supplementation did not affect body and muscle mass in aged male or female mice (Figure 1B-1D). This was consistent with no changes to organ mass or food intake arising from BAM15 supplementation (Supp. Figure 1).

**Figure 1.**
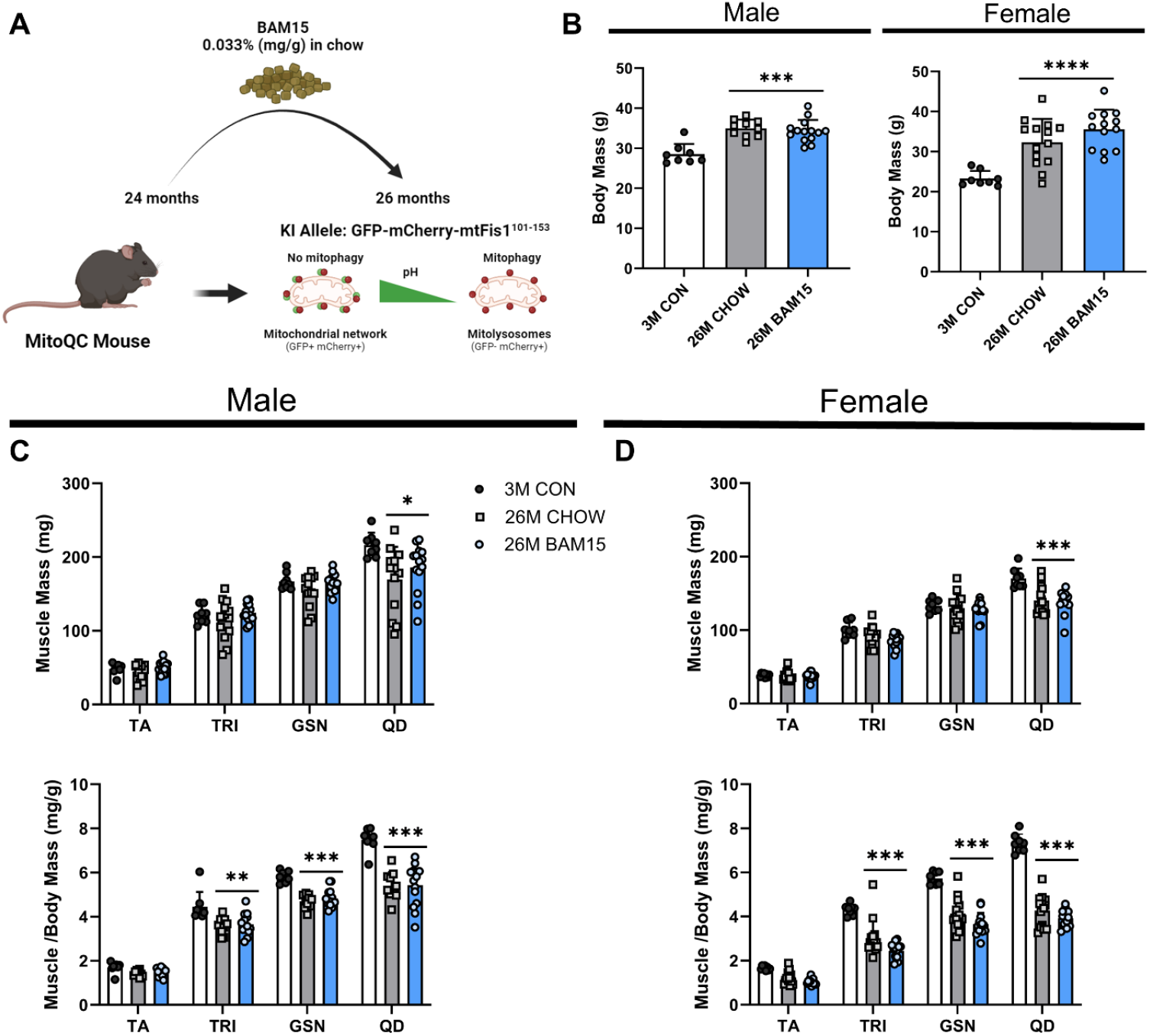
Body mass and muscle mass are unchanged from BAM15 supplementation in aged sarcopenic mice. A) Infographic of the mouse model and experimental design. B) Endpoint body mass, C) skeletal muscle mass (TA: tibialis anterior; TRI: triceps; GSN: gastrocnemius; QD: quadriceps) and D) skeletal muscle to body mass ratios shown. Data is presented as mean ± SD. n=7-8 per group for 3M CON; n=13-14 per group for 26M males and females. \**p*,0.05; \*\**p*<0.01; \*\*\**p*<0.001 vs 3M CON.

### BAM15 supplementation improves skeletal muscle contractile function in aged mice

In males, BAM15 supplementation protected against the age-associated loss of contractile function in EDL muscles, as represented by absolute and specific force frequency relationships similar to 3M CON (Figure 2A). BAM15 supplementation preserved EDL muscle characteristics relative to 3M CON, including; mass, optimal length, twitch force and the rate of force development, which were all reduced in the male 26M CHOW group (Supp. Table 1). BAM15 supplementation resulted in a greater resilience to fatigue in EDL muscles, with improved force production during the recovery period (Figure 2A). These findings were not reflected in male SOL data, which demonstrated lower absolute and specific force-frequency relationships, and reduced resilience to fatigue with lower force production during the recovery period in all aged groups (Figure 2B). However, SOL twitch characteristics demonstrated that BAM15 supplementation preserved time-to-peak, which was slower in the 26M CHOW group, suggesting potential protection to myosin-ATPase functionality without overall improvement in force production (Supp. Table 1). There were no additional effects on SOL twitch characteristics observed (Supp. Table 1). Consistent with male data, BAM15 supplementation protected against the loss of contractile function in female EDL muscles, represented by absolute and specific force frequency relationships (Figure 2C). However, there were no changes to female EDL twitch characteristics from BAM15 supplementation (Supp. Table 2). EDL muscles demonstrated an increase in the elasticity index, suggesting increased muscle stiffness in aged females (Supp. Table 2). BAM15 supplementation did not affect muscle resilience to fatigue in female EDL muscles (Figure 2C). Further, BAM15 did not affect contractility in female SOL muscles, with lower absolute and specific force-frequency relationships, and reduced resilience to fatigue with lower force production during the recovery period observed (Figure 2D). The rate of force development was lower in aged females (Supp. Table. 2), with no additional significant findings found for SOL twitch characteristics.

**Figure 2.**
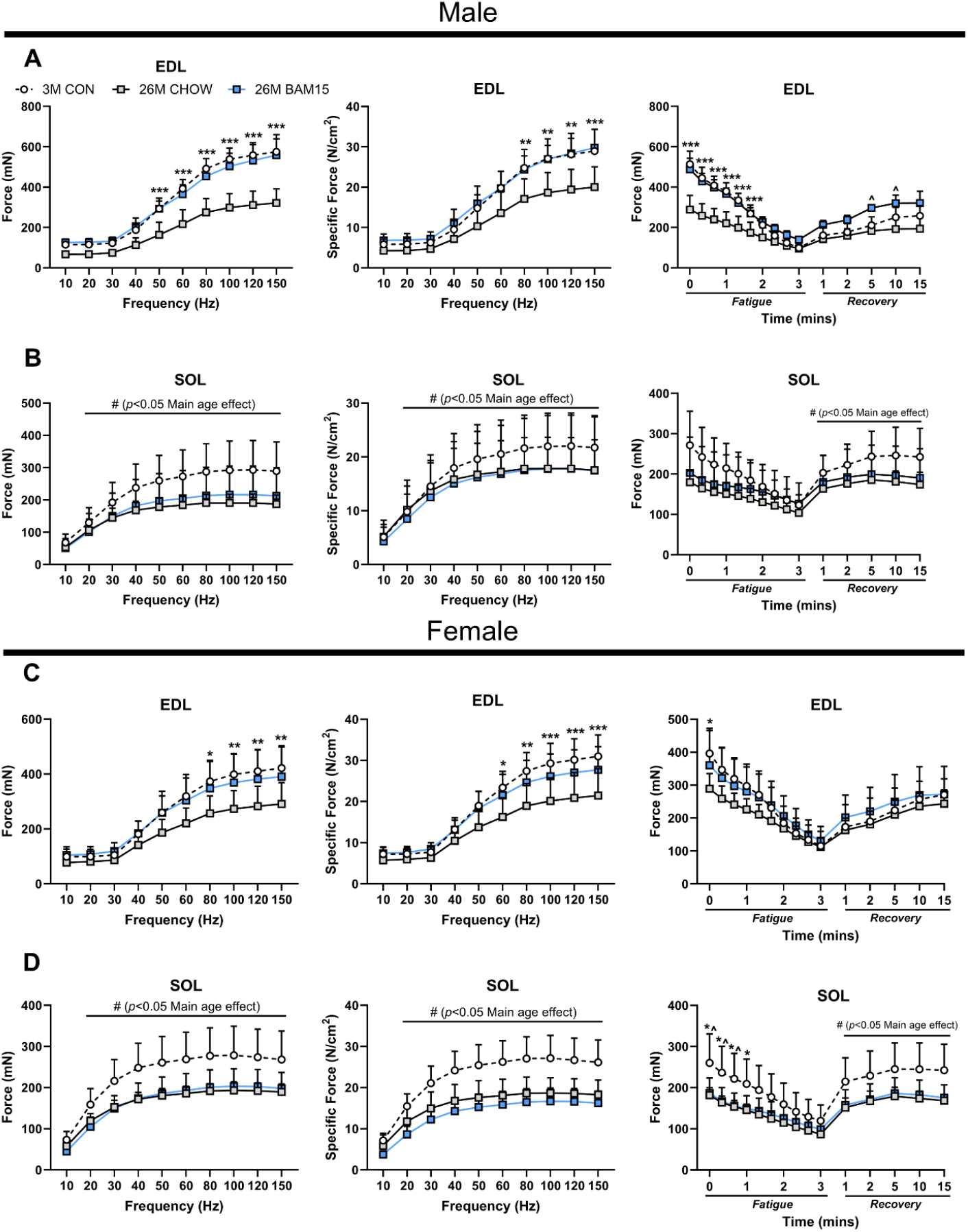
BAM15 supplementation improves EDL muscle contractile function in sarcopenic male and female mice. Ex vivo contractile function assessment was performed in extensor digitorum longus (EDL) and soleus (SOL) muscles. Male A) EDL and B) SOL muscles are analysed with force-frequency relationships showing absolute and specific force (normalised to physiological cross-sectional area) production across a range of frequencies in the EDL. Muscle resilience was tested with absolute force produced during the fatigue stimulus and recovery period shown. This contractile testing battery was repeated in female C) EDL and D) SOL muscles. Data is presented as mean ± SD. n=8 for 3M CON males and n=5-6 per group for 26M old males. n=8-9 per group for females. \**p*<0.05; \*\**p*<0.01; \*\*\**p*<0.001 26M CON vs 3M CON. ^ *p*<0.05 26M BAM15 vs 3M CON.

### BAM15 supplementation protects mitochondrial respiration in aged male mice

In permeabilised GTN muscle fibre bundles from males, BAM15 supplementation increased ADP-dependent respiration, but did not alter ADP sensitivity (Figure 3A). Apparent Km data suggest that the mitochondrial ‘fitness’ to respond to ADP as a substrate is near completely diminished in aged muscles (∼98% loss), due to enzyme active sites reaching saturation at low ADP concentrations (Fig. 3B). BAM15 supplementation protected against loss of Complex I leak respiration, suggestive of mild-uncoupling (Figure 3B). Further, BAM15 supplementation protected against age-related declines in Complex I and Complex I+II substrate-dependent respiration, and maximally uncoupled respiration in sarcopenic male muscles (Figure 3B). To investigate whether changes in mitochondrial respiration were connected to changes in mitochondrial content or oxidative stress, we measured mitochondrial ETC complex and citrate synthase protein content via immunoblotting. There were no changes to the protein content of mitochondrial ETC complexes I-V nor citrate synthase in aged male muscles (Figure 3C-D). However, BAM15 supplementation protected against elevated 4-HNE content (lipid-peroxidation marker) in aged male muscles (Figure 3C-D).

**Figure 3.**
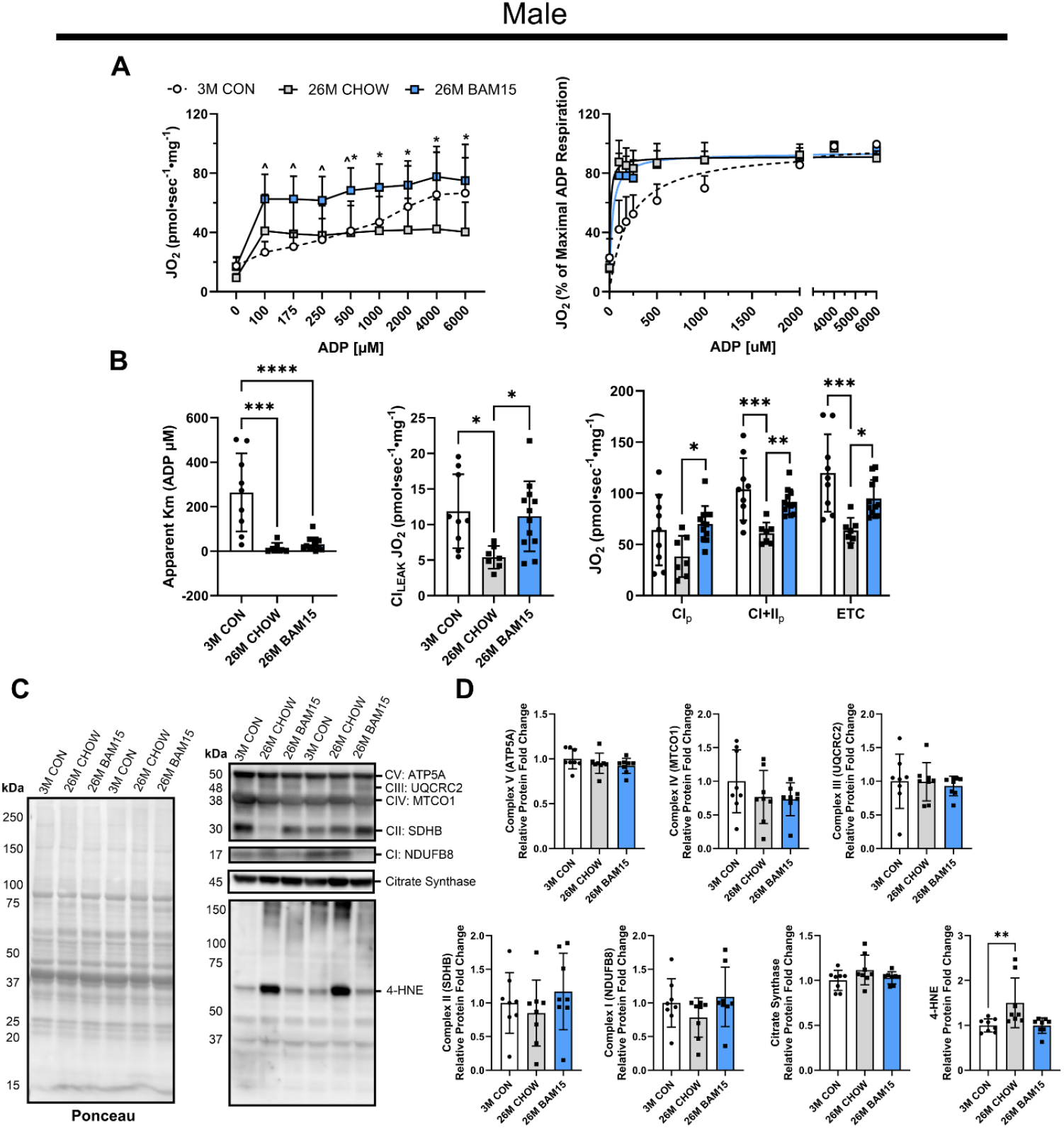
BAM15 supplementationpreserves skeletal musclemitochondrial respirationbut does not impact content abundance in sarcopenic male mice. Using permeabilized fibre bundles from male gastrocnemius muscles mitochondrial respiration was assessed via high resolution respirometry. A) Respiration is stimulated via an ADP titration in the presence of pyruvate and malate with standard and transformed data (via the Michaelis– Menten kinetics equation) presented. B) Apparent Km which represents the required concentration of ADP required to reach 50% of the maximal ADP-dependent respiration, Complex 1 Leak respiration, phosphorylating respiration (CI_p_) in the presence of Complex I substrate (glutamate) and Complex I+II substrates (glutamate and succinate – CI+II_p_) and CCCP-induced maximal uncoupled respiration are shown. Protein abundance was measured via immunoblotting with OXPHOS subunits Complex V (ATP5A), Complex IV (MTCO1), Complex III (UQCRC2), Complex II (SDHB) and Complex I (NDUFB8), alongside Citrate Synthase and 4 - Hydroxynonenal (4-HNE) probed. C) Representative images for antibodies and Ponceau for protein loading are shown and D) data quantified relative to 3M CON. Data is presented as mean ± SD. n=7-12 per group for mitochondrial respiration analyses; n= 7-8 per group for immunoblotting analyses. \**p*,0.05; \*\**p*<0.01; \*\*\**p*<0.001; \*\*\*\**p*<0.0001 26M CON vs 3M CON. ^ *p*<0.05 26M BAM15 vs 3M CON.

BAM15 supplementation did not protect against the reduction of ADP-dependent respiration and near-complete ablation of ADP sensitivity (∼97% loss) in aged female muscles (Figure 4A). We did not observe a loss in Complex I leak respiration in 26M CHOW group. Subsequently, there was no change in mitochondrial uncoupling observed through BAM15 supplementation (Figure 4B). Aged female muscles had lower Complex I and Complex I+II substrate-dependent respiration, and maximally uncoupled respiration compared to young reference controls, with no effect observed from BAM15 supplementation. Interestingly, we demonstrated a reduction in Complex I subunit NDUFB8 content in aged female muscles (∼30%), with some potential remodelling of Complex IV and Complex III content from BAM15 supplementation evident (Figure 4C-D). These findings may account, in part, for the diminished mitochondrial respiration. There were no further changes to Complex II and V, citrate synthase or 4-HNE content in aged females (Figure 4C-D).

**Figure 4.**
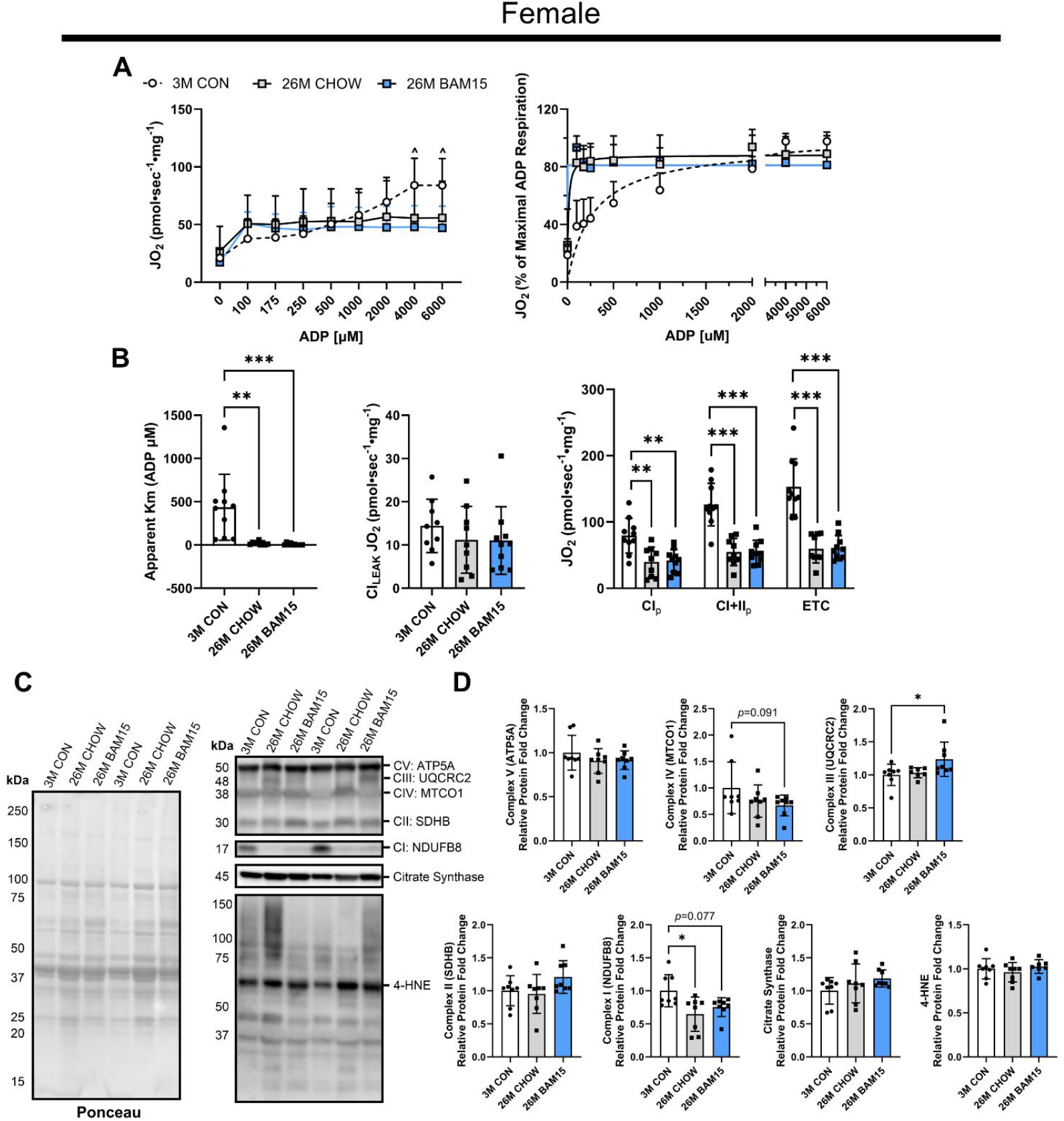
BAM15 supplementation does not alter skeletal muscle mitochondrial respiration or content abundance in sarcopenic female mice. Using permeabilized fibre bundles from male gastrocnemius muscles mitochondrial respiration was assessed via high resolution respirometry. A) Respiration is stimulated via an ADP titration in the presence of pyruvate and malate with standard and transformed data (via the Michaelis–Menten kinetics equation) presented. B) Apparent Km which represents the required concentration of ADP required to reach 50% of the maximal ADP-dependent respiration, Complex 1 Leak respiration, phosphorylating respiration (CI_p_) in the presence of Complex I substrate (glutamate) and Complex I+II substrates (glutamate and succinate – CI+II_p_) and CCCP-induced maximal uncoupled respiration are shown. Protein abundance was measured via immunoblottingwith OXPHOS subunits Complex V (ATP5A), Complex IV (MTCO1), Complex III (UQCRC2), Complex II (SDHB) and Complex I (NDUFB8), alongside Citrate Synthase and 4 -Hydroxynonenal (4-HNE) probed. C) Representative imagesfor antibodies and Ponceau for proteinloadingare shown and D) data quantified relative to 3M CON. Data is presented as mean ± SD. n=9-10 per group for mitochondrial respiration analyses; n= 7-8 per group for immunoblotting analyses. \**p*,0.05; \*\**p*<0.01; \*\*\**p*<0.001; \*\*\*\**p*<0.0001 26M CON vs 3M CON. ^ *p*<0.05 26M BAM15 vs 3M CON.

### BAM15 supplementation does not alter skeletal muscle mitophagy in aged mice

BAM15 administration has previously been suggested to alter mitophagy-related protein content in a mouse model of sarcopenic-obesity [10]. Thus, we employed the use of MitoQC mice to directly test whether BAM15 alters skeletal muscle mitophagy. In aged male mice, we demonstrate there are no changes to mitolysosome abundance from BAM15 supplementation (Fig. 5A-C). When normalised to fibre area, there was no effect of age or BAM15 supplementation observed on skeletal muscle mitophagy (Fig. 5D) However, we did observe an increase in mitolysosome size in male 26M muscles (∼52%, Fig. 5E). This may indicate alterations in mitophagic processes or machinery which drive the accumulation of larger mitolysosomes. In aged female mice we observed an increase in mitolysosome abundance (∼76%) which is not rescued by BAM15 supplementation (Fig. 6A-C). This is compatible with normalised data, with unsignificant trends indicating an increase in mitolysosome per fibre area (Fig. 6D). Consistent with aged male data, female 26M mice demonstrated an increase in mitolysosome size (∼49%, Fig. 6E).

**Figure 5.**
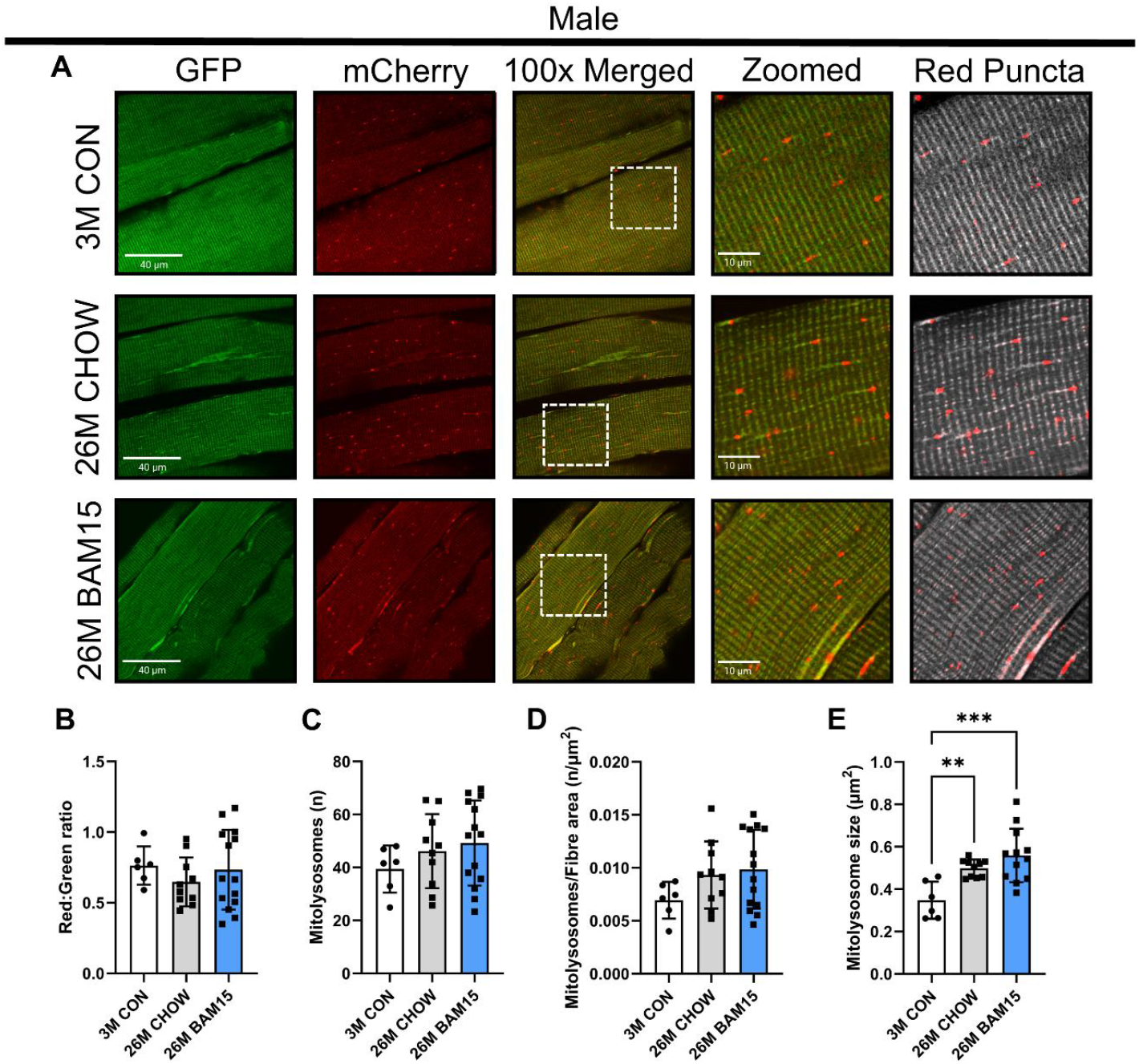
BAM15 supplementation does not alter skeletal muscle mitophagystatus insarcopenic malemice. Fixed quadriceps muscles were embedded in OCT and longitudinally sectioned for mitophagy quantitation via measurement of MitoQC reporter fluorophores GFP and mCherry using confocal microscopy. A) Representative images are displayed, alongside B) mean intensity ratio of the red and green channels, C) total number of mitolysosomes, D) mitolysosomes per fibre area and E) mitolysosome size reported. Data is presented as mean ± SD. 15-17 fibres analysed per section. n=6 for 3M CON; n= 10-14 per group for 26M males. \**p*,0.05; \*\**p*<0.01; \*\*\**p*<0.001; 26M CON vs 3M CON.

**Figure 6.**
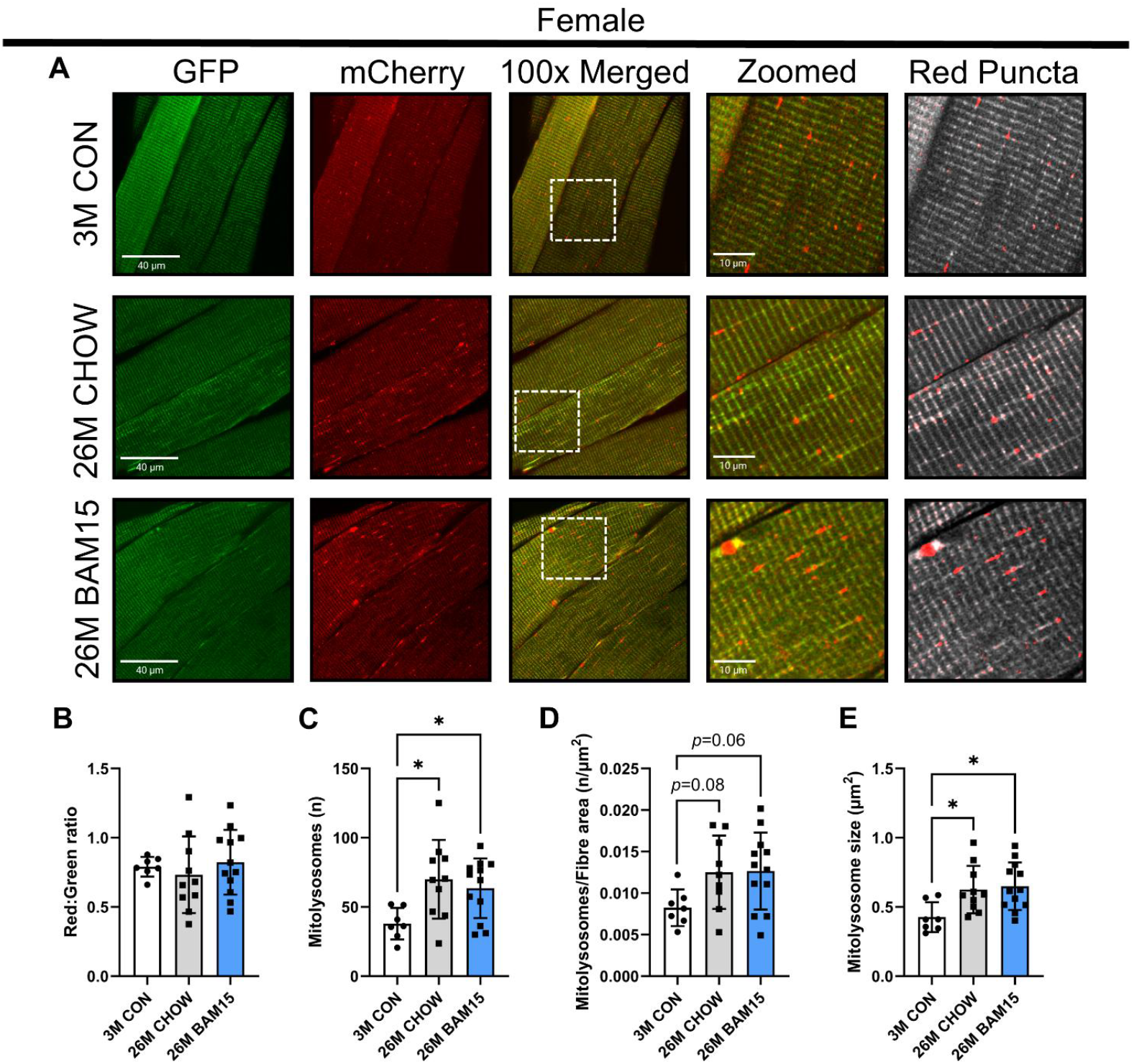
BAM15 supplementation does not alter skeletal muscle mitophagy status in sarcopenic female mice. Fixed quadriceps muscles were embedded in OCT and longitudinally sectioned for mitophagy quantitation via measurement of MitoQC reporter fluorophores GFP and mCherry using confocal microscopy. A) Representative images are displayed, alongside B) mean intensity ratio of the red and green channels, C) total number of mitolysosomes, D) mitolysosomes per fibre area and E) mitolysosome size reported. Data is presented as mean ± SD. 15-17 fibres analysed per section. n=7 for 3M CON; n= 10-12 per group for 26M females. \**p*,0.05; \*\**p*<0.01; \*\*\**p*<0.001; 26M CON vs 3M CON.

## Discussion

Here we provide the first evaluation of the mitochondrial uncoupler BAM15 as a sarcopenia drug candidate in male and female mice. We demonstrate that BAM15 supplementation does not alter age-associated loss in muscle mass, but prevents a decline in skeletal muscle contractility and mitochondrial respiration leading to functional improvements in sarcopenic muscle. Protection against age-related muscle functional decline was evident in both male and female EDL muscle force production. This was consistent with previous investigation by Dantas *et al*. where BAM15 supplementation improved 4-limb grip strength in aged high fat diet-induced obese mice [10]. Functional protection from BAM15 supplementation in aged muscle may be explained, in part, by the preservation of muscle bioenergetics, observed as improved mitochondrial respiration and the restoration of oxidative stress marker 4-HNE. Oxidative stress has been shown to negatively impact excitation-contraction coupling through impaired Ca^2+^ sensitivity in aged mice and mice with global deletion of CuZn superoxide dismutase, that have lower antioxidant activity [19]. This mechanism linking preserved muscle bioenergetics is likely to indirectly involve protection against motor neuron degeneration, which is required for skeletal muscle dysfunction during sarcopenia [20].

In aged male mice, BAM15 supplementation enhanced skeletal muscle resilience to fatigue. This is likely related to improved mitochondrial function but is difficult to definitively pinpoint, given the complex nature of skeletal muscle fatigue [21]. In aged females, BAM15 supplementation did not show changes to fatigue resilience. We suggest this is due to female aged EDLs having a lower delta change in contractile force production due to age when compared to males (∼107mN lower in female and ∼241mN lower in male aged controls compared to young references). In EDL muscle, there seems to be a clear relationship between muscle force production and muscle resilience to fatigue which may be a better indicator of overall muscle performance rather than fatigue susceptibility. Age-related muscle decline has a greater burden in predominantly fast type II myo-fibre muscles [1], which aligns with our data in the predominantly type IIB myo-fibre EDL muscle. Despite being susceptible to age-related functional decline, SOL muscle contractility data did not follow the same protective theme arising from BAM15 supplementation. There are examples of paradoxical mechanistic regulation between EDL and SOL muscles, such as global myostatin knockout mice, which exhibit lower specific force production in EDL muscles, while SOL muscles remain unaffected compared to wild-type controls [22]. A simpler explanation could be that SOL (mixed myo-fibre type) muscle does not undergo mitochondrial decline to the same extent as the EDL, (predominately type II myo-fibre type) during aging [23].

Previous work has demonstrated that the mitochondrial-targeted antioxidant mitoquinone mesylate (mitoQ) did not maintain EDL muscle force production of aged mice [24], whilst the mitochondrial-targeting peptide SS-31 (elamipretide) improved treadmill running performance and reduced fatigue susceptibility without changes to force production in tibialis anterior muscles of aged mice [25, 26]. Further, delivery of untargeted, mito-protective compounds such as NAD^+^ precursors [27, 28], oleuropein [29] and Urolithin A [30], demonstrate improved functional capacity and/or muscle contractility in aged mice. BAM15 therefore joins a short list of mitochondrial targeted compounds that can improve sarcopenic muscle function.

BAM15 supplementation preserved substrate-dependent phosphorylating respiration and maximally uncoupled ETC-linked respiration without any changes in mitochondrial protein content in aged male mice. To date, *in vivo* data has only shown that BAM15 supplemented in high-fat diet can increase mitochondrial content and Complex II-IV activity in skeletal muscles of aged male mice [10]. In our hands, BAM15 supplementation protected mitochondrial function, which is underscored by the attenuation of the oxidative stress marker, 4-HNE. Tsuji *et al*. previously demonstrated that BAM15 improves survival in a mouse model of sepsis by protecting kidneys from oxidative damage and lowering circulating mtDNA [31]. These findings are linked to the mild uncoupling induced by BAM15, a targeted mechanism that has long been considered to reduce oxidative/nitrosative stress and restore normative mitochondrial function [6, 32]. This will also lead to lower ROS production rate and protection against unchecked oxidative damage from reactions instigated by single electron reductants that can react with free oxygen and lead to superoxide (O_2_^-^) formation [7]. Mild uncoupling induced by BAM15 supplementation is therefore a promising anti-oxidant strategy to protect tissues from oxidative damage and mitochondrial respiration.

Skeletal muscle from aged female mice demonstrated a different mitochondrial phenotype compared to males. Like male counterparts, aged female mice had lower substrate-dependent phosphorylating and maximally-uncoupled respiration. However, this was not underlined by changes in oxidative stress or mitochondrial content markers, rather a ∼30% loss of Complex I subunit NDUFB8 protein content. To the best of our knowledge, this finding has not previously been observed. Recently, data from Moreira-Pais *et al*. suggested that the age-dependent remodelling of the skeletal muscle mitochondrial proteome is more pronounced in females due to reduced serum 17β-estradiol levels and the mitochondrial pool of estrogen receptor alpha (ERα) [33]. These authors also identified downregulation of Complex 1 subunit NDUFV2 from proteomic analysis [33], suggesting some overlap between our findings. In aged female muscles we also observe higher rates of mitophagy, with no changes in males.

Loss of ADP sensitivity is considered to be a key representation of the metabolic decline in aged human skeletal muscle, which is a reversible bioenergetic target of resistance training [34]. However, our data show that aged mouse muscles show complete loss of the metabolic fitness to respond to ADP as a substrate, with near-complete saturation observed after 100 μM of ADP, regardless if supplemented with BAM15. Pharaoh *et al*. found that there is also an increase in the amount of ADP required to reduce H_2_O_2_ production in aged mouse muscle [35]. This was underlined by lower content of proteins involved in the transport of ADP into mitochondria, ADP/ATP translocase 2 (ANT2) and the voltage-dependent anion channel (VDAC) [35]. Additionally, there was no change in mitophagy levels from BAM15 supplementation in aged muscles. However, there was some indication of altered mitophagic processes, with larger mitolysosomes identified in both aged male and female muscles [36, 37]. Collectively, these data suggest that there is likely a nuanced redox-dependent mechanism that regulates the relative loss of metabolic fitness to respond to ADP as a substrate and ADP transporter abundance, which in turn can alter mitophagy in aged muscles.

In summary, BAM15 supplementation preserved EDL contractile function and mitochondrial respiration without altering age-related loss in muscle mass. Future investigations should explore both earlier and extended treatment durations for BAM15 supplementation, given that age-related muscle decline is typically considered to commence at ∼18 months of age in mice. Our findings suggest that maintaining mitochondrial efficiency and redox balance may mitigate functional decline in sarcopenic muscle. Together, our data support mitochondrial uncoupling as a promising therapeutic strategy to preserve muscle function during the progression of sarcopenia.

## Acknowledgements

This work was supported by funding from the Australian National Health and Medical Research Council awarded to A.P. (APP2013278). I.A. and A.P. were supported by Wellcome Leap’s Dynamic Resilience Program (jointly funded by Temasek Trust).

This research was facilitated by access to Sydney Mass Spectrometry, a core research facility at the University of Sydney. The authors acknowledge the technical and scientific assistance of Sydney Microscopy & Microanalysis, the University of Sydney node of Microscopy Australia.

K.L.H. and W.L.S. declare commercial interest in Life Biosciences Inc and Uncoupler Biosciences.

