## Supplementary Data for "Mitochondrial uncoupler BAM15 improves skeletal muscle function and mitochondrial respiration in Sarcopenia"

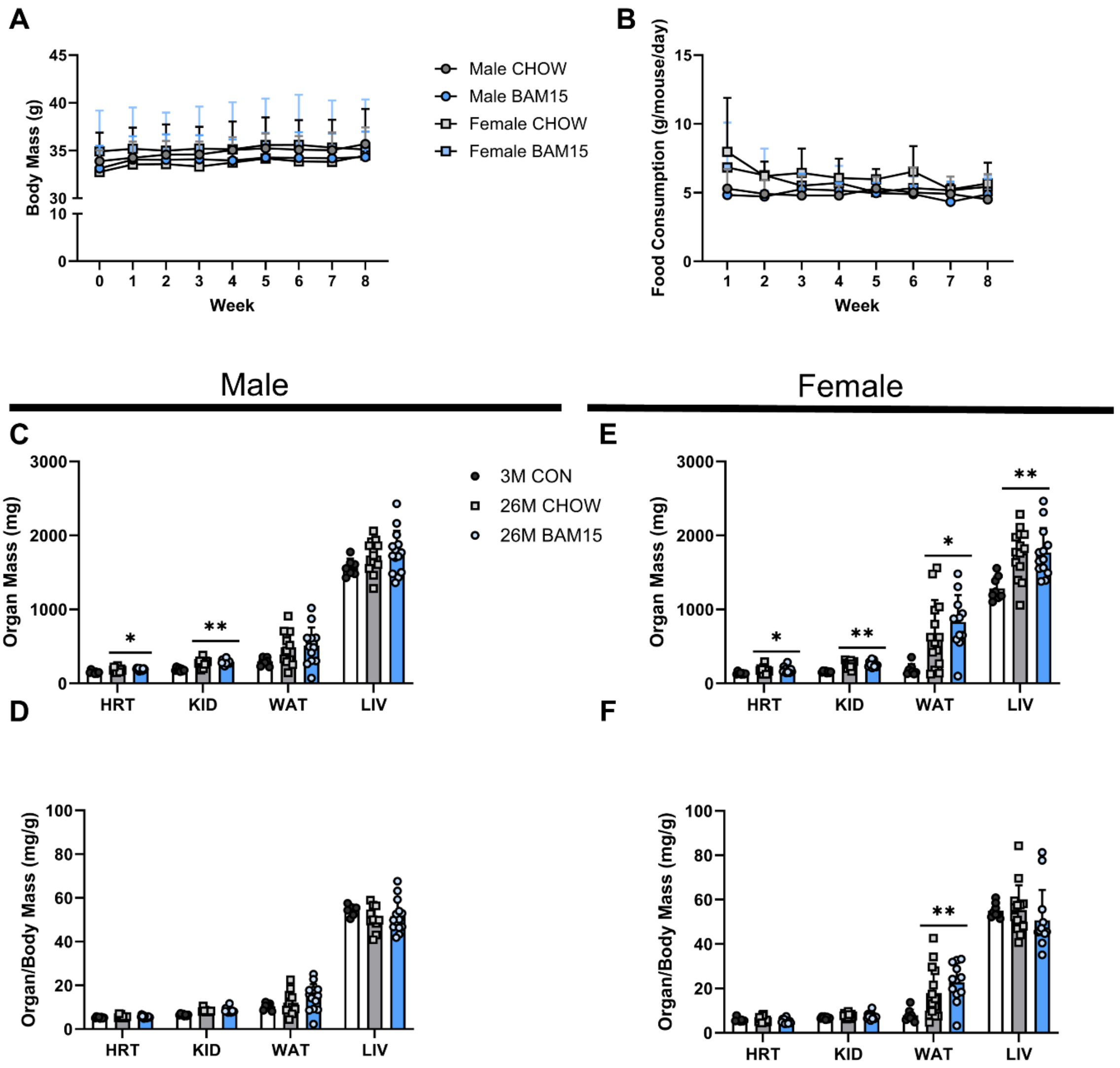


**Figure 1: Body mass, food intake and endpoint organ mass are unchanged from BAM15 supplementation in aged sarcopenic mice.** A) Body mass and B) food intake is displayed over the 8-week diet intervention. C) organ mass (HRT: heart; KID: kidney; WAT: white adipose tissue from epidydimal fat pad; LIV: liver) and D) organ to body mass ratios are shown. Data is presented as mean ± SD. n=7-8 per group for 3M CON; n=13-14 per group for 26M males and females. **p*,0.05; ***p*<0.01; ****p*<0.001 vs 3M CON.

**Supplemental Table 1: Detailed male ex vivo skeletal muscle contractile properties analyses.** Lo: Optimal Length; pCSA: Physiological cross-sectional area; Po: Absolute force; sPo: Specific Force; Pt: Twitch Force; TTP: Time to twitch peak; 1/2RT: Half-relaxation time (twitch). Data is presented as mean ± SD. n= 7-8 per group for 3M CON, n=5-6 per group for 26M groups.

| | **EDL** | | | | |
|---|---|---|---|---|---|
| | **3M CON** | **26M CHOW** | **26M BAM15** | **p value (3M CON vs 26M CHOW)** | **p value (3M CON vs 26M BAM15)** |
| **Mass (mg)** | 12.1 ± 1.2 | 9.6 ± 1.6 | 11.3 ± 1.6 | 0.0134* | 0.5706 |
| **Lo (mm)** | 13.3 ± 0.4 | 12.3 ± 0.8 | 12.9 ± 1.1 | 0.0460* | 0.4902 |
| **pCSA (cm$^2$)** | 0.019 ± 0.002 | 0.017 ± 0.003 | 0.019 ± 0.002 | 0.1507 | 0.9683 |
| **Po (mN)** | 575.1 ± 64.8 | 334.2 ± 69.7 | 558.6 ± 101.6 | <0.0001*** | 0.9077 |
| **sPo (N/cm$^2$)** | 30.6 ± 3.0 | 20.1 ± 4.5 | 29.8 ± 4.6 | 0.0005*** | 0.9239 |
| **Pt (mN)** | 95.3 ± 17.8 | 65.9 ± 11.0 | 113.4 ± 14.5 | 0.0047** | 0.0958 |
| **TTP (s)** | 20.4 ± 1.5 | 19.7 ± 0.6 | 20.9 ± 1.5 | 0.4985 | 0.7292 |
| **1/2RT (s)** | 11.0 ± 1.4 | 12.7 ± 1.0 | 9.9 ± 1.1 | 0.0802 | 0.3927 |
| **Rate of force development (mN/s)** | 4.7 ± 0.7 | 3.3 ± 0.5 | 5.5 ± 1.1 | 0.0095** | 0.1266 |
| **Elasticity Index (Pt/Po)** | 0.17 ± 0.04 | 0.20 ± 0.03 | 0.21 ± 0.04 | 0.1701 | 0.1166 |
| | **SOL** | | | | |
| | **3M CON** | **26M CHOW** | **26M BAM15** | **p value (3M CON vs 26M CHOW)** | **p value (3M CON vs 26M BAM15)** |
| **Mass (mg)** | 10.2 ± 1.1 | 7.9 ± 1.1 | 9.1 ± 0.9 | 0.0020* | 0.1656 |
| **Lo (mm)** | 10.3 ± 0.7 | 9.3 ± 1.1 | 9.5 ± 1.2 | 0.1514 | 0.2686 |
| **pCSA (cm$^2$)** | 0.013 ± 0.002 | 0.011 ± 0.002 | 0.013 ± 0.002 | 0.1769 | 0.9580 |
| **Po (mN)** | 293.4 ± 90.6 | 190.6 ± 25.0 | 217.3 ± 87.9 | 0.0433* | 0.1474 |
| **sPo (N/cm$^2$)** | 22.1 ± 5.7 | 17.5 ± 5.3 | 17.8 ± 10.3 | 0.3584 | 0.1105 |
| **Pt (mN)** | 47.0 ± 17.1 | 32.1 ± 6.8 | 35.5 ± 19.2 | 0.1686 | 0.3200 |
| **TTP (s)** | 29.6 ± 1.8 | 32.4 ± 1.9 | 28.9 ± 2.1 | 0.0288* | 0.7743 |
| **1/2RT (s)** | 28.7 ± 3.2 | 31.3 ± 4.9 | 25.9 ± 3.5 | 0.3828 | 0.3218 |
| **Rate of force development (mN/s)** | 1.6 ± 0.6 | 1.0 ± 0.2 | 1.2 ± 0.7 | 0.2007 | 0.4405 |
| **Elasticity Index (Pt/Po)** | 0.17 ± 0.07 | 0.17 ± 0.02 | 0.16 ± 0.03 | 0.9992 | 0.8977 |

**Supplemental Table 2: Detailed female ex vivo skeletal muscle contractile properties analyses.** Lo: Optimal Length; pCSA: Physiological cross-sectional area; Po: Absolute force; sPo: Specific Force; Pt: Twitch Force; TTP: Time to twitch peak; 1/2RT: Half-relaxation time (twitch). Data is presented as mean ± SD. n= 7-8 per group for 3M CON, n=8-9 per group for 26M groups.

| | **EDL** | | | | |
|---|---|---|---|---|---|
| | **3M CON** | **26M CHOW** | **26M BAM15** | **p value (3M CON vs 26M CHOW)** | **p value (3M CON vs 26M BAM15)** |
| **Mass (mg)** | 7.8 ± 0.8 | 8.3 ± 0.7 | 8.5 ± 0.9 | 0.3654 | 0.1542 |
| **Lo (mm)** | 12.3 ± 0.7 | 13.1 ± 0.6 | 13.1 ± 0.6 | 0.0430* | 0.0275* |
| **pCSA ($cm^2$)** | 0.014 ± 0.001 | 0.014 ± 0.001 | 0.014 ± 0.002 | 0.9925 | 0.8307 |
| **Po (mN)** | 421.8 ± 80.8 | 313.6 ± 47.9 | 390.7 ± 108.0 | 0.0423* | 0.6784 |
| **sPo ($N/cm^2$)** | 31.0 ± 5.2 | 23.2 ± 4.7 | 27.7 ± 5.6 | 0.0246* | >0.9999 |
| **Pt (mN)** | 87.6 ± 24.9 | 86.0 ± 21.5 | 93.6 ± 28.5 | 0.9893 | 0.8451 |
| **TTP (s)** | 20.7 ± 1.0 | 21.0 ± 0.6 | 21.1 ± 1.9 | 0.9134 | 0.7617 |
| **1/2RT (s)** | 12.2 ± 1.1 | 12.7 ± 1.7 | 13.0 ± 1.9 | 0.7672 | 0.4699 |
| **Rate of force development (mN/s)** | 4.2 ± 1.2 | 4.1 ± 1.0 | 4.5 ± 1.5 | 0.9699 | 0.8413 |
| **Elasticity Index (Pt/Po)** | 0.21 ± 0.04 | 0.24 ± 0.03 | 0.24 ± 0.03 | 0.0405* | 0.0580 |
| | **SOL** | | | | |
| | **3M CON** | **26M CHOW** | **26M BAM15** | **p value (3M CON vs 26M CHOW)** | **p value (3M CON vs 26M BAM15)** |
| **Mass (mg)** | 7.4 ± 1.1 | 7.4 ± 1.8 | 8.4 ± 1.0 | 0.9988 | 0.3142 |
| **Lo (mm)** | 9.6 ± 1.0 | 9.1 ± 0.7 | 9.1 ± 1.2 | 0.5436 | 0.5255 |
| **pCSA ($cm^2$)** | 0.011 ± 0.002 | 0.011 ± 0.003 | 0.012 ± 0.002 | 0.8200 | 0.1590 |
| **Po (mN)** | 279.8 ± 69.7 | 195.4 ± 18.7 | 203.8 ± 41.8 | 0.0072** | 0.0149* |
| **sPo ($N/cm^2$)** | 27.3 ± 5.4 | 18.9 ± 3.8 | 16.7 ± 3.2 | 0.0026** | 0.0003*** |
| **Pt (mN)** | 42.5 ± 13.2 | 33.0 ± 5.3 | 28.6 ± 5.9 | 0.1106 | 0.0165* |
| **TTP (s)** | 31.5 ± 1.4 | 33.6 ± 3.0 | 30.7 ± 2.6 | 0.1870 | 0.7246 |
| **1/2RT (s)** | 32.6 ± 2.4 | 33.8 ± 5.2 | 30.9 ± 3.2 | 0.7613 | 0.5786 |
| **Rate of force development (mN/s)** | 1.4 ± 0.4 | 0.98 ± 0.13 | 0.93 ± 0.17 | 0.0421* | 0.0202* |
| **Elasticity Index (Pt/Po)** | 0.152 ± 0.03 | 0.171 ± 0.03 | 0.143 ± 0.03 | 0.3923 | 0.8072 |
